# An ultrasensitive T-cell receptor detection method for TCR-Seq and RNA-Seq data

**DOI:** 10.1101/740340

**Authors:** Si-Yi Chen, Qiong Zhang, Chun-Jie Liu, An-Yuan Guo

## Abstract

T-cell receptors (TCRs) recognizing antigens play vital roles in T-cell immunology. Surveying TCR repertoires by characterizing complementarity-determining region 3 (CDR3) can provide valuable insights into the immune community underlying pathologic conditions, which will benefit neoantigen discovery and cancer immunotherapy. Here we present a novel tool named CATT, which can apply on TCR sequencing (TCR-Seq), RNA-Seq, and single-cell TCR(RNA)-Seq data to characterize CDR3 repertoires. CATT integrated maximum-network-flow based micro-assembly algorithm, data-driven error correction model, and Bayes classification algorithm, to self-adaptively and ultra-sensitively characterize CDR3 repertoires with high accuracy. Benchmark results of datasets from *in silico* and real conditions demonstrated that CATT showed superior recall and precision compared with other prevalent tools, especially for datasets with short read length and small data size. By applying CATT on a TCR-Seq dataset from aplastic anemia patients, we found the skewing of TCR repertoire was due to the oligoclonal expansion of effector memory T-cells. CATT will be a powerful tool for researchers conducting TCR and immune repertoire studies. CATT is freely available at http://bioinfo.life.hust.edu.cn/CATT.

## Introduction

T-cells as important immune effector cells play important roles in cell-mediated immunity and cancer immunotherapy. T-cell receptors (TCRs) are disulfide-linked, membrane-anchored, heterodimeric proteins located on the surface of T-cells, and function in adaptive immune response by recognizing specific antigens (Medzhitov and Janeway 1997). Investigation of TCR repertoires by TCR profiling can provide prized views for understanding the functions of T-cell in immune processes (e.g., immune responses, immunosuppression, and immunotherapies, Kessels et al. 2001; Pogorelyy et al. 2019; Schuster et al. 2011). For example, TCR profiling has been used to monitor the status of T-cells underlying disease progression (Attaf et al. 2015) and drug therapies (Thommen et al. 2018), which can assist in early-stage diagnosis (de Masson et al. 2018) and precision medicine (Ott et al. 2017). Additionally, characterization of TCR repertoires of tumor-infiltrating T-cells can benefit the selection of cancer treatments (Zacharakis et al. 2018) and prognostic prediction (Page et al. 2016), while identification of neoantigen-reactive T-cell clones by TCR profiling is the most critical step that directly impact the curative effect of TCR gene therapy (Harris and Kranz 2016). The diversity and antigen-specificity of TCRs are mainly determined by complementarity-determining region 3 (CDR3), which is the most hypervariable region generated by V(D)J recombination (Clambey et al. 2014). Thus, surveying CDR3 repertoires can, therefore, offer qualitative insights into TCR repertoire and immunological research (Turner et al. 2006).

High-throughput sequencing, including both bulk and single-cell TCR/RNA sequencing (scTCR-Seq/scRNA-Seq), has provided advanced conveniences for investigating CDR3 repertoires (Brown et al. 2015). However, due to the natures of CDR3s and limitations of technologies (*e.g.*, the highly diverse and numerous rare clonotypes for CDR3s, errors induced by PCR and sequencing), accurate characterization of CDR3 repertoires remains a great challenge. To date, several tools, such as MiXCR (Bolotin et al. 2017), RTCR (Gerritsen et al. 2016), IMSEQ (Kuchenbecker et al. 2015), LymAnalyzer (Yu et al. 2016), TraCeR (Stubbington et al. 2016), and TRUST (Hu et al. 2019), have been developed for to characterizing CDR3 repertoires in TCR-Seq and/or RNA-Seq data. However, these methods have some limitations and require improvements in terms of 1) discarding reads without complete CDR3s or with low frequency, which may cause massive loss of CDR3 sequences and bias toward TCR repertoires, 2) parameter sensitivity and reliance on hands-on settings for model optimization, 3) poor performance in datasets with short read length and low TCR content, and 4) excessive consumption of time and computational resource, which roadblocks the usage of these methods

To overcome these limitations, we developed a sensitive and accurate method, named CATT (***C***har***A***cterizing ***T***CR reper***T***oires, http://bioinfo.life.hust.edu.cn/CATT), for characterizing CDR3 repertoires in both bulk and single-cell TCR(RNA)-Seq datasets. CATT showed superior accuracy and sensitivity compared with other five popular tools (LymAnalyzer, RTCR, IMSEQ, TRUST, and MiXCR) on *in silico* and real datasets, and could provide more unbiased information of TCR repertoires in immune response and immunotherapy.

## Results

### Overview and main steps of CATT

For better characterization of CDR3 repertoires on bulk and single cell TCR(RNA)-Seq data, the algorithm designed by CATT comprises three main steps: 1) *De novo* assembly of CDR3 sequences using reads aligned to the loci of known TCRs by maximum-network-flow algorithm, 2) Error correction for CDR3 sequences by a data-driven model balanced the base quality and reads coverage, and 3) Confidence assessment and annotation for CDR3 sequences by a Bayesian statistical method. The overview of the core algorithm of CATT is shown in Fig 1, and the detailed illustration is in the Materials and Methods section. The benchmark of CATT and other five popular tools (LymAnalyzer, RTCR, IMSEQ, TRUST, and MiXCR) in curated *in silico* and real datasets is presented in the following sections.

**Figure 1.**
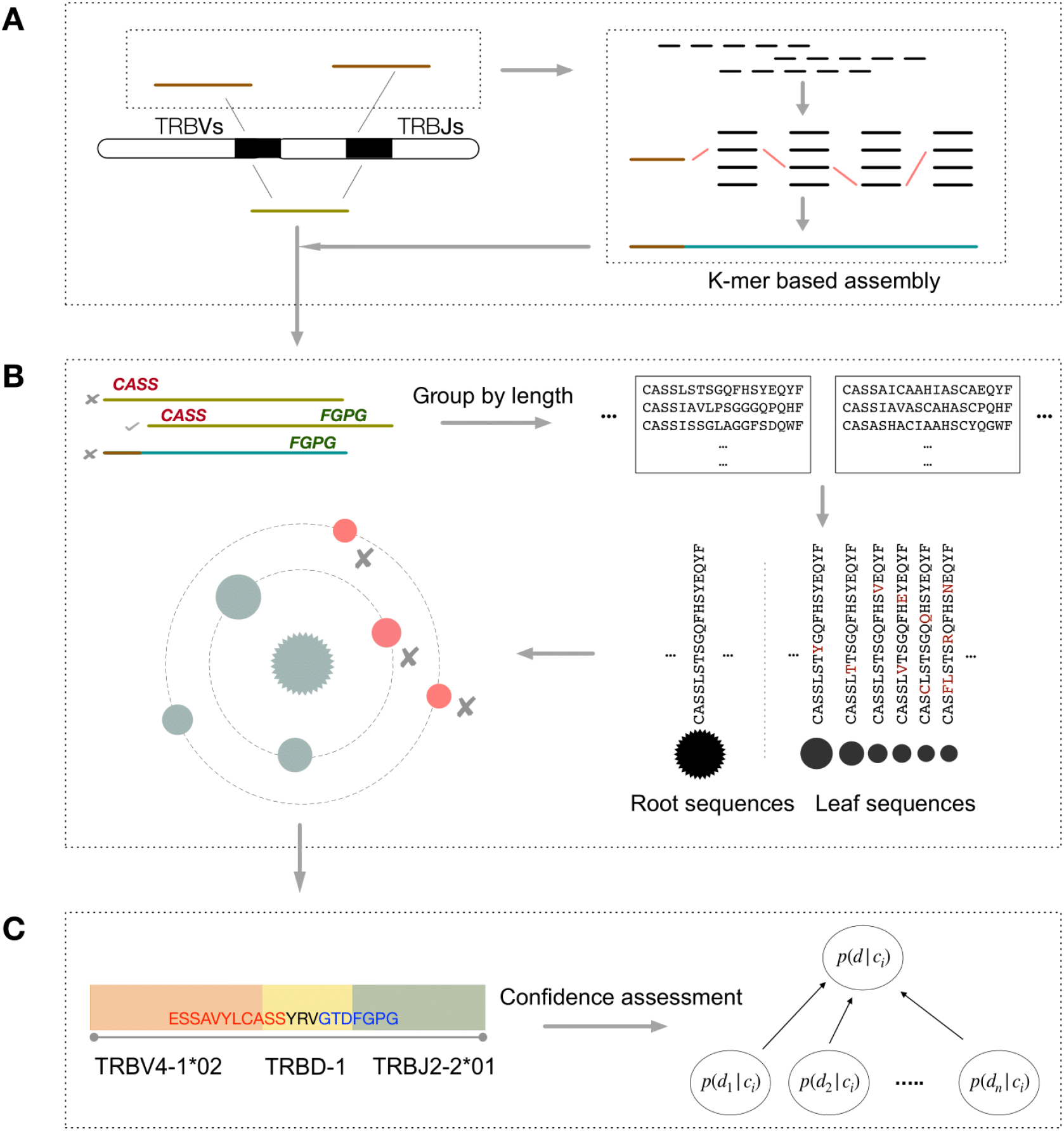
Overview of the core algorithm of CATT. (A) Candidate CDR3 detection. All reads are aligned to V and J reference genes to select out candidate (brown) reads for micro-assembly. Potential CDR3 sequences were reconstruct by k-1 overlapped k-mers

### Performance of CATT in *in silico* datasets

To better mimic *in silico* TCR-Seq and RNA-Seq data, several factors that could impact the efficiency of tools in real situations, such as the amplification error from PCR, sequencing error, read length, data size, and TCR content, were introduced in our simulation (details are provided in the “Method” section). Four prevalent tools (MiXCR, IMSEQ, RTCR, and LymAnalyzer) were included to benchmark the performance in TCR-Seq datasets, whereas TRUST was supplemented with the 4 tools in RNA-Seq dataset (TRUST is only limited in RNA-Seq). Performance was evaluated at two levels: (1) clonotype, measured by the recall and precision of detected CDR3s, and (2) repertoire, which considers the difference in CDR3 distribution between *in silico* datasets and outcomes of each tool, as measured by deviations in distribution calculated using Kullback-Leibler divergence (KLD) (Kullback and Leibler 1951).

The performance of CATT exhibited competitive advantages in the *in silico* TCR-Seq datasets compared with other tools (Fig. 2A, B), especially in datasets with shorter read lengths (Fig. 2A). For the clonotype level, the recall of CATT showed almost 2–3 times higher than other tools in datasets with read lengths of 75 and 100 bp, while the precision of CATT maintained at a high and robust level (mean±SD, 93.52±1.01%). Meanwhile, on the repertoire level, CATT also showed a lower deviation of CDR3 distribution between the *in silico* datasets and outcomes compared with other tools, suggesting that the CDR3 repertoires characterized by CATT were closer to real situations. Additionally, we observed that the deviation of CDR3 distribution showed a decreasing tendency with increases in data size and read length (Fig. 2B), indicating the accuracy of characterizing CDR3 repertoires can be improved with more data capacity and a longer read length. Overall, CATT robustly recovered most CDR3 sequences in all datasets, with a high sensitivity and precision at both clonotype and repertoire levels.

**Figure 2.**
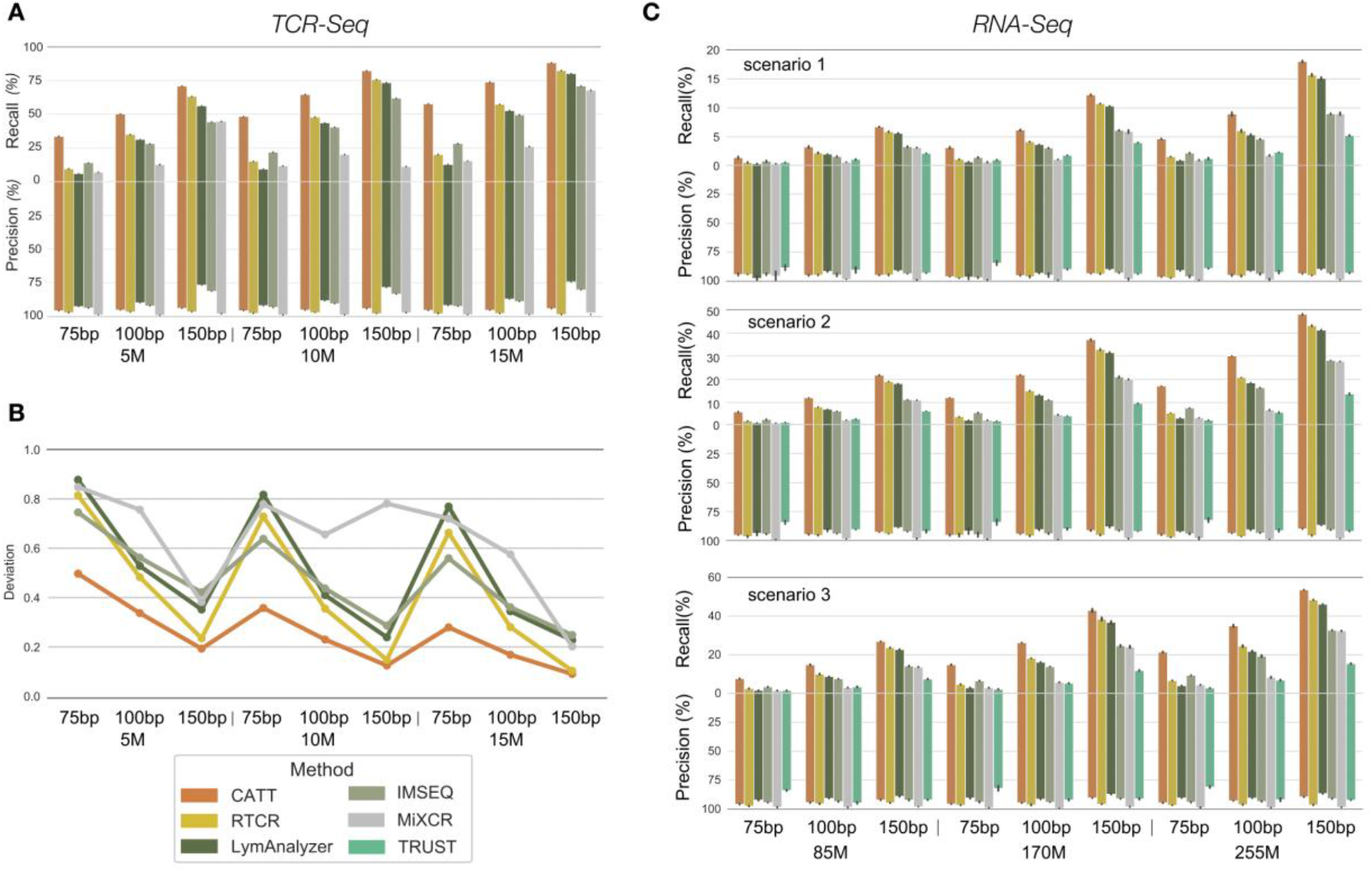
Performance of CATT and other tools in in silico data. (A) The recall and precision of each tool for CDR3 sequences in in silico TCR-Seq datasets. (B) The distribution deviation of CDR3 repertoires for each tool in in silico TCR-Seq datasets. (C) The recall and precision of each tool for CDR3 sequences in in silico RNA-Seq datasets.

Besides the evaluation of CATT in TCR-Seq data, we also assessed the performance of CATT in *in silico* RNA-Seq data, which contain three RNA-Seq datasets for the simulation of different tissues. Due to the low content of TCRs in RNA-Seq data, the overall recall of each tool declined in comparison with the results in TCR-Seq data (Fig. 2C). However, CATT still showed an advanced sensitivity and precision in all the RNA-Seq datasets compared with other tools. In particular, the recall of CATT was almost two times higher than that of other tools in datasets with read lengths of 75 and 100 bp, and was approximately 20% higher when the read length was 150 bp. Although the precision of most tools fluctuated with the data size, CATT showed a stable and high precision in all datasets (mean±SD, 93.78±1.75%), suggesting that CATT performed well in RNA-Seq data.

In addition, we evaluated the computational consumption (*i.e.*, memory usage and time cost) for CATT in the *in silico* datasets (Supplemental Fig. S1). CATT consumed acceptable memory usage and relatively low time costs in all the datasets, while the time and memory costs of CATT were slowly increased with data size and read length.

### Performance of CATT in real datasets

To assess the performance of CATT in real situations, we further applied it in four published datasets: 1) scTCR-Seq dataset of patients with basal cell carcinoma (NCBI BioProject: PRJNA509910; 34 samples), 2) scRNA-Seq dataset of CD4^+^ T_reg_ and T_mem_ cells from different human tissues (Patil et al. 2018) (NCBI BioProject: PRJEB22806; 2037 samples), 3) bulk TCR-Seq with unique molecular identifiers (UMI) labeled dataset of patients with neurological immune-mediated disorders (Alves Sousa et al. 2019) (NCBI BioProject: PRJNA495603; 106 samples), and 4) bulk RNA-Seq dataset of patients with melanoma with paired TCR-Seq data (Bolotin et al. 2017) (BioProject: PRJNA371303; 2 samples).

Due to lack of the ground truth of CDR3 repertoires in real datasets, detected CDR3 with the following features were considered as high-confident ones used in this evaluation: 1) For single-cell TCR-Seq or RNA-Seq datasets, the high-confident CDR3s were from majority voting results of each tool; 2) For the bulk UMI-labeled TCR-Seq datasets, the high-confident CDR3 sequences were supported by more than 3 UMIs and spanning the entire reference CDR3 region; 3) For bulk RNA-Seq datasets, detected CDR3s recurring from paired TCR-Seq data were acted as high-confident ones. The detailed procedures were in the “Materials and Method” section.

Considering each T-cell usually generates one kind of TCR (Brady et al. 2010), the scTCR-Seq data is an ideal resource for assessing the performance of tools in CDR3 detection. Similar to the performance in *in silico* datasets, CATT exhibited great advantages compared with the other tools in both scTCR-Seq and scRNA-Seq datasets (Fig. 3A, B). For example, CATT showed a high and robust recall as well as high precision in scTCR-Seq dataset (the left one of Fig. 3A and Supplemental Fig. S2), while CATT detected the least number of CDR3 clonotypes in most cells (Fig. 3A, right), which appeared to be closer to real situations. The number of detected CDR3 clonotypes varied greatly among different tools in the scTCR-Seq data of each cell (Fig. 3A, right), suggesting severe false positive errors and large deviations between real situations and outcomes of tools. Besides, CATT successfully detected CDR3s in 77.2% of samples in scRNA-Seq datasets, which was almost 15% higher than that with other tools (Fig. 3B). Most of CDR3s detected by CATT were reoccurring in the outcomes of other tools, suggesting that CATT achieved a high precision in scRNA-Seq data as well.

**Figure 3.**
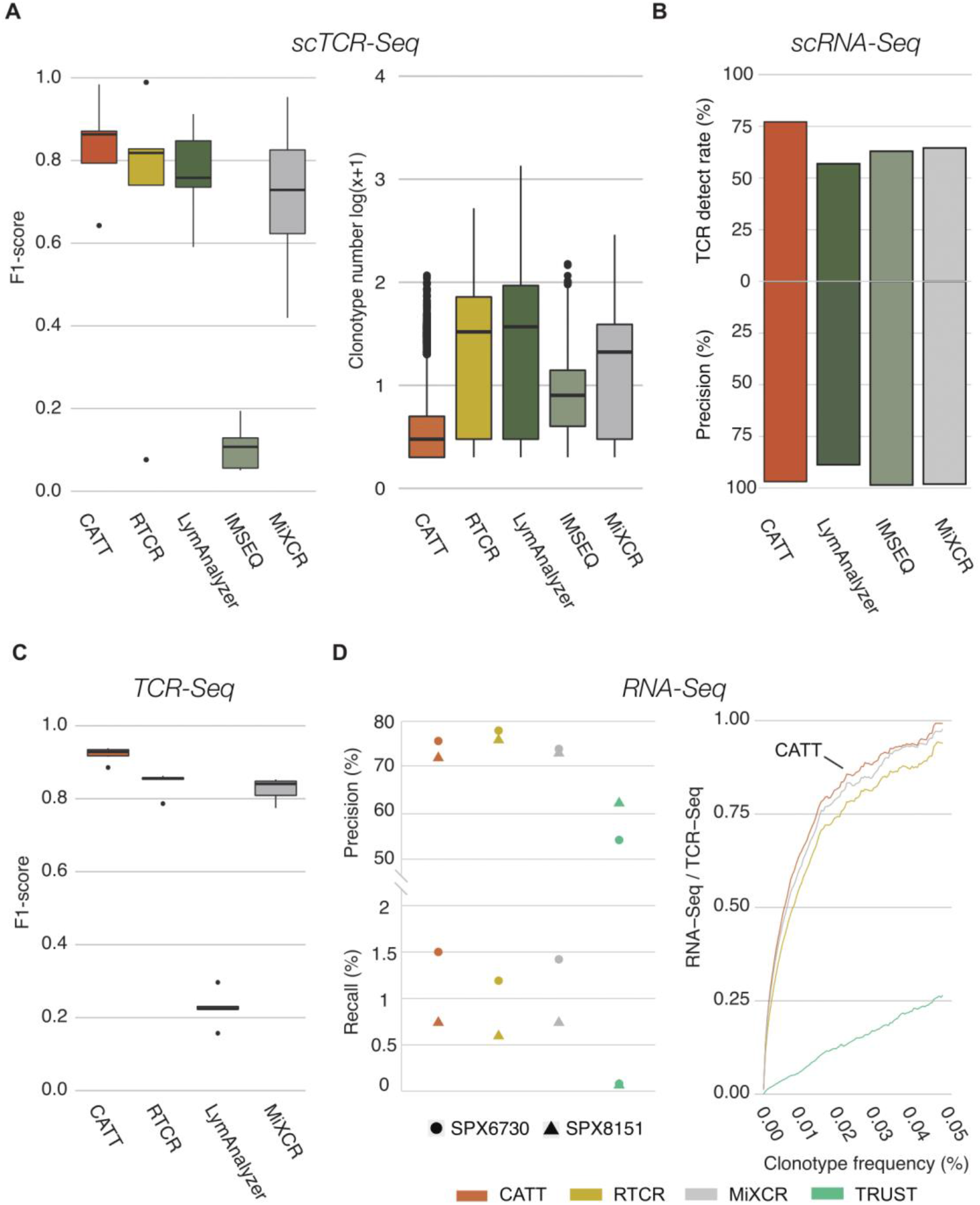
Performance of CATT and other tools in real datasets, including scTCR-Seq, scRNA-Seq, TCR-Seq, and RNA-Seq data. (A) The F1-score and detected CDR3 clonotype of each tool in scTCR-Seq datasets. F1-score is in the left, while the right one is the numbers of detected CDR3 clonotype in each cell. (B) The precision and recall of each tool in scRNA-Seq datasets. (C) The precision and recall of each tool in real TCR-Seq datasets. (D) The precision and recall of each tool in real RNA-Seq datasets (left one), while the right curve displays the consistency of CDR3 repertoires from paired RNA-Seq and TCR-Seq datasets.

In evaluations for real bulk TCR-Seq datasets, CATT demonstrated superior performance (Fig. 3C) with the highest precision (Supplemental Fig. S3). Although all tools exhibited a similar recall on TCR-Seq datasets (~95%), the precision of CATT was remarkably higher than that of other tools. Due to the low TCR content in RNA-Seq data, though the recall of all tools fallen at a low level, CATT showed the highest recall as well as high precision among all tools, indicating that CATT is suitable for the application on real data (Fig. 3D, left). Moreover, the outcomes of CATT from paired RNA-Seq and TCR-Seq datasets showed higher coincidence compared with other tools (Fig. 3D, right), implying that CATT could detect CDR3 clonotypes with high sensitivity. Taken together, CATT exhibited a robust efficiency (high sensitivity and precision) on characterizing CDR3 repertoires in the TCR-Seq, RNA-Seq, and single-cell sequencing datasets. Some tools had no output in the particular dataset (IMSEQ for TCR-Seq, LymAnalzyer in RNA-Seq, etc.), which may because of the tools require higher TCR content, sequence length and data quality.

### A case study of CATT identified TRBV skewing in CD8^+^ T cell population by the oligoclonal expansion of effector memory T cells (T_EM_)

CDR3 repertoires can provide important diagnostic biomarkers and serve as a surrogate predictor for diseases and prognosis (McNeel 2016). To reveal the biological meanings of CDR3 repertoires detected by CATT, we employed a TCR-Seq data of patients with aplastic anemia (GSE101660, 42 samples) as a case study. Aplastic anemia is an autoimmune disease characterized by the destruction of hematopoietic progenitor or stem cells (Ishiyama 2016), in which the abnormal TCR signaling may play pivotal roles in the process (Xiao et al. 2017). The aplastic anemia dataset comprises 24 samples (12 for CD8^+^ T cells, 10 for CD4^+^ T cells, and 2 for CD8^+^CD57^+^ T cells) from 12 patients and 18 samples (8 for CD8^+^ T cells and 10 for CD4^+^ T cells) from 9 healthy donors.

In CD8^+^ T cells, the length distribution of CDR3 repertoires in healthy donors was close to the normal distribution, with a peak at 15 AA, whereas the peak switched to14 AA in patient samples (Fig. 4A). No significant difference was observed between CD4^+^ T cells between healthy donors and patients (Fig. 4A). Additionally, the usage of TRBV genes in CD8^+^ T cells differed between patients and healthy donors, but this phenomenon was not observed in CD4^+^ T cells (Supplemental Fig. S4). Taken together, the shorter length of CDR3 sequences and skewed usage of TRBV genes suggests an aberrance of CD8^+^ T cells in patients with aplastic anemia.

**Figure 4.**
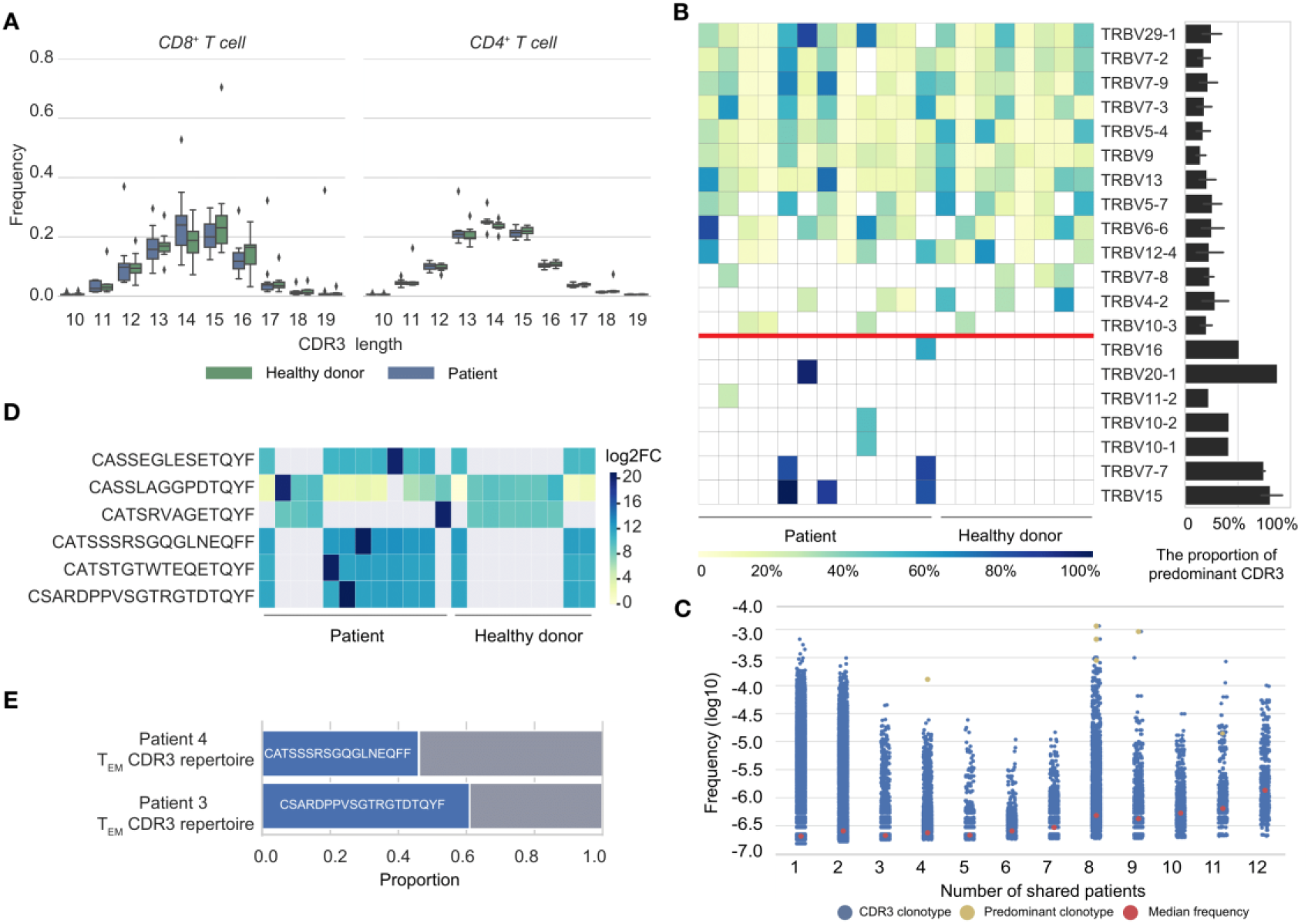
A case study of CATT on CDR3 characterization from aplastic anemia dataset. (A) The length distributions of CDR3 sequences in the CD4+ and CD8+ T cells. (B) The usage of TRBV gene (heatmap) and the frequency of predominant CDR3s in the TRBV gene (bar plot). The TRBV genes below the red line were patient-specific. (C) The frequencies of shared CDR3s across patients. Each blue point represents a CDR3 clonotype, yellow points represent the predominant CDR3s in patient-specific TRBV genes, and the red dots represent the median frequency of CDR3s shared by a corresponding number of samples. (D) The relative deviation of predominant CDR3s between patients and healthy donor (patients-VS-healthy donors). (E) The ratio of predominant CDR3s in the CDR3 repertoires of TEM cells from two patients.

In all the prevalent TRBV genes (top 10 of each sample) used in CD8^+^ T cells, several TRBV genes were patient-specific that not appeared in healthy donors (Fig. 4B). The predominant CDR3s were found oligoclonally expanded in the CDR3 repertoires of these patient-specific TRBV genes, (Fig. 4B). Meanwhile, these clonotypes were widespread in patients with a relatively high frequency (Fig. 4C) and functioned as the predominant clonotype in the entire CDR3 repertoire (Fig. 4D). Furthermore, among these CDR3 clonotypes, “CATSSSRSGQGLNEQFF” and “CASRDPPVSGTRGTFTQYF” were found in the TCR-Seq data of CD8^+^ CD57^+^ T cells (effector memory T cells, T_EM_), accounting for approximately 50% of the T_EM_ CDR3 repertoire in the two patients (Fig. 4E). The results were consistent with the flow cytometry findings of original publication that the oligoclonal expansion of T_EM_ is observed in the development of aplastic anemia (Giudice et al. 2018). Combined with the results of the performance of CATT in different datasets and its real-world application, CATT has the potential to be the preferred tool to investigate CDR3 repertoires.

## Discussion

T-cells as important components of the adaptive immune system play vital roles in host defense, and the interaction between TCRs and antigens is a key step for the functions of T-cells (Wu et al. 2014). The enormous diversity and dynamics of TCR repertoires are mainly determined by hypervariable CDR3 polymorphisms, and the current common strategy for TCR repertoire profiling is CDR3 characterization (Heather et al. 2018). In this study, we developed a computational method, CATT, which employs a novel assembly and an error correction model to characterize CDR3 repertoires in both bulk and single-cell TCR(RNA)-Seq data.

CDR3 characterization provides opportunities for interpretation of immune repertoires, vaccine profiling, neoantigen discovery, and the development of immunotherapy (Ott et al. 2017; Roth et al. 2018). The precise characterization of CDR3 repertoires is the most critical and basic issue in the associated field. Although a few methods have been developed for CDR3 characterization, several limitations roadblock their wide applications. For example, TRUST (Hu et al. 2019) was specifically designed to detect CDR3 sequences in bulk RNA-Seq data; LymAnalyzer (Yu et al. 2016), RTCR (Gerritsen et al. 2016), and IMSEQ (Kuchenbecker et al. 2015) rely on reads covered the entire CDR3 region or directly mapped to V(D)J genes, ignoring reads partially aligned to CDR3 regions. However, these reads can provide an imperative clue for CDR3 communities, particularly for rare clonotypes. Therefore, discarding these reads will result in the loss of information and falsification of clonotype frequency. Moreover, although IMSEQ and MiXCR (Brady et al. 2010) employ model-based sequence correctors to remove rare clonotypes, they appear to be parameter sensitive and require appropriate parameters for different datasets. CATT employs a new and effective strategy to detect CDR3 sequences. The high sensitivity and precision of CATT are mainly due to the maximum-network-flow and data-driven-transition-probability learning algorithms. Reads partially aligned to the V or J region were used to construct a k-mer dictionary, then the maximum-network-flow algorithm was employed to assign these k-mers to optimal graphs and assemble potential CDR3s based on the k-mer frequency. The data-driven-transition-probability learning algorithm was designed to eliminate erroneous clonotypes, which calculates the confidence probability for each CDR3 using binomial distribution method to prevent absorbing erroneous or candidate-error rare clonotypes.

CATT exhibited robustly high sensitivity and accuracy both in the TCR-Seq and RNA-Seq data, especially in the single cell TCR(RNA)-Seq data (Fig. 2A, C, 3A). These characteristics can provide comprehensive views for investigating the features and functions of T-cells in specific biological processes, such as carcinogenesis and immunotherapy (Ellsworth et al. 2017; Zhang et al. 2016; Zheng et al. 2017; Abdelmoez et al. 2018). Although the low abundance of TCRs in bulk and single cell RNA-Seq data may result in the incomplete estimation of TCR repertoires, CATT achieved a high-quality performance as well (Fig. 2C). Considering the monoclonal condition for CDR3 repertoires within a given scTCR-Seq dataset, the bias of outcomes of CATT was significantly smaller than other tools (Fig. 3C). The superior sensitivity of CATT can provide more benefits in the detection of rare CDR3s and modeling of the dynamics of TCR repertoires, and can help discerning TCR recognition or designing therapeutic TCR molecules (Leisegang et al. 2016; Morris and Stauss 2016; Hinrichs et al. 2017). Besides, the high sensitivity of CATT can reduce the data size demands and cost barriers. However, due to the hypervariability of CDR3 and limitations of the algorithm designed by CATT, unmapped reads that may contain additional information to recover entire CDR3 repertoires were discarded. Indeed, TCR recognition event may not require the complete matching between CDR3 sequences and antigens (Glanville et al. 2017). Moreover, entire CDR3 sequences can provide more comprehensive insights into the features and functions of T-cells underlying the interplays between antigens and immune system.

Existing tools for CDR3 characterization mainly rely on reads that cover approximately full-length regions of CDR3 and require targeted and complete amplification of CDR3 loci during experimental steps. CATT could characterize CDR3 repertoires with high accuracy and predominant sensitivity, even for single-cell sequencing datasets. Compared with the other prevalent tools in different datasets, CATT offers prior advantages of CDR3 repertoire characterization with sensitivity, accuracy, and high-resolution profiling. CATT can serve as a preferred tool for the characterization of TCR repertoires and the discovery of candidate biomarkers, which will benefit personalized TCR T-cell therapy and neoantigen-specific T-cell discovery.

## Materials and Methods

### Algorithm designed by CATT

Briefly, CATT detects CDR3 sequences by employing reads aligned to the loci of known TCRs and *de novo* reassembling these reads using a feasible maximum-network-flow algorithm. Reads with both ends spanning the V and J genes contained candidate CDR3 sequences; while reads partially mapped to the V or J genes were used for assembly. After assembling, CATT employed the motif pattern of known CDR3s from the IMGT (Lefranc et al. 2015) resource to measure whether the candidate and assembled CDR3 sequences were putative CDR3s. Next, CATT merged ultralow-frequency putative CDR3 sequences with the high-frequency ones using a data-driven model. Finally, CATT assessed the confidence of putative CDR3 sequences using a supervised Bayesian classification method. Details of the algorithm and procedures are described below.

#### Detection and assembly for CDR3 sequences

CATT aligned reads to VJ reference genes by bowtie2 (Langmead and Salzberg 2012) with the sensitive model. Reads mapped to both V and J genes were deemed to comprise the entire CDR3 (named CDR3 reads), whereas reads marked as soft-clipped or partially aligned to the V or J genes were kept for a further assembly (named candidate reads). Firstly, a k-mer frequency table was built using the candidate reads, and k-mers from the 3′ end of candidate reads that mapped to V genes (named as leader kmers) were used as templates to construct potential CDR3 sequences (k − 1 overlap stepping to J genes or vice versa). While none overlap k-mer used for further extension or the length of assembled sequences reached 150 bp, all the selected k-mers formed feasible flows, and the path of each flow represented candidate CDR3 sequences. Next, the frequency of the leader kmer was used as a source, and the maximum-network-flow algorithm was employed to assign the source to each flow passed by recursive overlapping k-mers, whose combinations can carve-up the source approximately.

#### Pattern match

The motif pattern from IMGT criteria (CDR3 sequences started with amino acid C at 104 and ended with a FGXG motif) were used to measure the reliability of candidate and assembly CDR3 sequences. CATT enumerated all possible open reading frames (ORFs) for the above CDR3 sequences. CDR3 AA sequences satisfied the IMGT criteria and without any stop codon were kept for further analyses. Furthermore, CDR3 AA sequences with lengths in the range of [5, 35] were considered as putative CDR3s.

#### Error correction

CATT designed a novel data-driven algorithm to correct the biases from PCR amplification and sequencing errors, which considered the read length, the abundance of CDR3s, and sequences similarity among CDR3s. First, putative CDR3 sequences were divided into subgroups by their length (L). Subsequently, we hypothesized that erroneous CDR3 sequences stemmed from root sequences and were similar to root sequences but with a low frequency (Yokota et al. 2017). In each subgroup, sequences with a frequency lower than H were deemed as leaf sequences, whereas the other sequences were considered as root sequences. The parameter H was calculated using the following formula (Equation 1), which represents the maximum frequency of an erroneous sequence that was produced by a single base error:

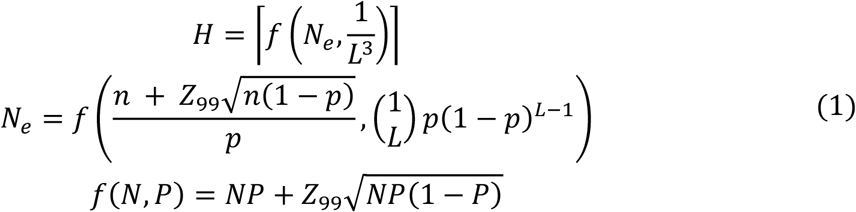

where *n* is the maximum frequency of CDR3 sequences in the corresponding subgroup, the parameter *L* represents the length of the CDR3 sequence with maximum frequency, *p* stands for the error rate of bases retrieved from the alignment results, *N*_*e*_ is the estimated frequency of all sequences with 1 error base.(from PCR or sequencing error), and *f*(*N*, *P*) represents the 99% upper confidence bound of the binomial distribution.

In each subgroup, the leaf sequences with Hamming distance less than or equal to *D* for a root sequence were considered as candidate erroneous sequences derived from the root one. The parameter *D* is the integer that:

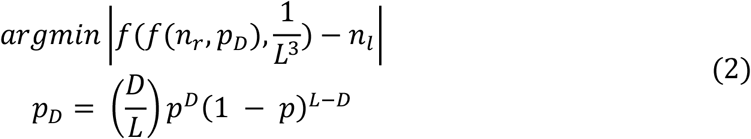

Where *p*_*D*_ denotes the probability of D error base in the root sequence, *n*_*r*_ and *n*_*l*_ represent frequencies of the root and leaf sequences, respectively, and *D* represents the maximum possible Hamming distance between the root and leaf sequences according to binomial distribution and their frequency. If the Hamming distance between the leaf and root sequences was higher than *D*, the leaf sequence is unlikely stemmed from the root sequence.

After the above process, CATT calculated the transition rate *S*_*j*_ from the root sequence to each paired leaf sequence to distinguish whether the leaf sequence stemmed from a sequencing error or a true event. The transition rate *S*_*j*_ is calculated as follows (Equation 3):

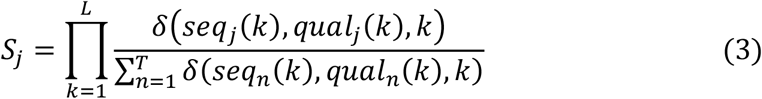

where *T* is the total number of leaf sequences; *seq*_*j*_(*k*) and *qual*_*j*_(*k*) represent the nucleotide type and quality score in the kth position of the leaf sequence j, respectively; and *δ*(*seq*_*j*_(*k*), *qual*_*j*_(*k*), *k*) are the total number of combination nucleotide *seq*_*j*_(*k*) and quality score *qual*_*j*_(*k*) in the kth position for all leaf sequences, respectively. The transition rate *S*_*j*_ denotes the possibility that all of the leaf sequences were derived from the different base replacement of the root sequence in the subgroup. CATT then merged the frequency of leaf sequences into that of the paired root sequence in ascending order of their frequency divided by the transition rate. To prevent rare CDR3 sequences from above procedures, the frequency of the root sequence after error correction was limited less than 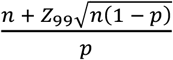 according to binomial distribution.

#### Annotation and confidence evaluation

After error correction, CATT realigned putative CDR3 sequences to TCR references to identify the usage of V, D, and J segment, and CDR3 sequences without V and J segments were discarded. Additionally, CATT employed a Bayes classifier to assess the confidence of CDR3 sequences as follows (Equation 4):

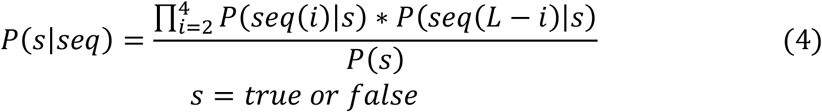

where *seq*(*i*) represents the ith nucleotide in the sequence and *L* is the sequence length. The training set for the classifier was downloaded from VDJdb (Shugay et al. 2018), which comprises 17,792 human TRB CDR3 sequences.

### *In silico* dataset preparation

A pool comprising 10^6^ synthetic TCR sequences was simulated according to the model presented by Murugan et al (Murugan et al. 2012). For TCR-Seq datasets, TCR sequences were randomly selected from the pool and assigned with corresponding abundances to mimic the heavy-tailed distribution of TCR sequences in real situations, which satisfied the Zipf distribution with the parameter of α = 3 (Bolkhovskaya et al. 2014). To simulate the errors introduced by PCR amplification in the library preparation procedure of TCR-Seq, selected TCR sequences were amplified approximately 18 times, with a reaction rate of *m* − 1 and a substitution error rate of 2 × 10^−5^ (Shagin et al. 2017), where *m* follows a normal distribution (E = 1.90, D = 0.1) (Karlen et al. 2007). ART (Huang et al. 2012) was then used to simulate paired-end reads of Illumina HiSeq 2500 (default error profiles provided by ART) on DNA libraries (mean fragment length 300 bp, standard deviation 100 bp) with different read lengths (75, 100, and 150 bp) and different library sizes (5, 10, and 15 M reads).

Due to various infiltration levels of lymphocytes in different tissues and RNA-Seq background noise, we simulated three RNA-Seq datasets for the liver, testis, and brain tissues. For each tissue, we used Polyster (Frazee et al. 2015) to generate simulated reads with different data sizes (85, 170, and 255 M) using corresponding transcriptome profiling from the GTEx project as a background. We then mixed TCR reads according to lymphocyte infiltration rates from the TCGA biospecimen data. T-cell infiltration in these tissues was 10% (the liver), 20% (the testis), and 50% (the brain), which represents the content of TCR reads at 4×10^−5^, 8×10^−5^, and 2×10^−4^, respectively (Brown et al. 2015). Considering the low abundance of TCRs in a real situation, we simulated a new pool of synthetic TCR sequences using the same process but with a Zipf distribution parameter of α = 2.5 and full-length TRBC gene segments.

### Strategies for assessing performance

TRUST (v3.0.2), IMSEQ (v1.1.0), RTCR (v0.4.3), MiXCR (v3.0.5), and LymAnalyzer (v1.2.2) were applied with default settings. For IMSEQ, “clustering-based error correction” and “merging of identical CDR3 sequences with ambiguous segment identification” were turned on, i.e., “-ma -qc -sc.” For TRUST, a STAR aligner (Dobin et al. 2013) was employed to align reads to the hg19 reference. For MiXCR, we added the “rna-seq” and “assemblePartial” parameters for RNA-Seq datasets. All methods were performed with 16 threads, and run times were measured on Ubuntu 12.04 system with 256 GB of RAM and Inter Xeon E7-4820.

The performance evaluation of each tool on different data was according to the following three criteria

1. *Clonotype level*. Measured by recall and precision. In *in silico* data, the output CDR3 sequences of each tool were compared with ground-true CDR3s to access the performance. As for real data, which lacked ground-true CDR3s, we distinguished “true” and “false” sequences based on the following rules:

1.1) For single-cell TCR(RNA)-Seq datasets, the true CDR3 of each cell was determined on the basis of majority voting results of all the tools. In cases of none CDR3 sequence or alternative CDR3 sequence detected in a single cell, the cell will be excluded from further analyses. Additionally, a true CDR3 sequence from scRNA-Seq data should comprise both alpha and beta information.
1.2) For bulk TCR-Seq datasets, CDR3 sequences supported by at least three UMIs and containing entire TRBV and TRBJ segment are considered as “true” ones.
1.3) For bulk RNA-Seq datasets, the true CDR3 sequences are the ones detected in the paired TCR-Seq data.
2. *Repertoire level:* Measured by the deviation (Equation 5) of distributions between the characterized CDR3 repertoires of tools and real situations. The F1 score (Equation 6) and Kullback–Leibler divergence (KLD, also known as relative entropy, Equation 7) were used to calculate the deviation (Kullback and Leibler 1951).

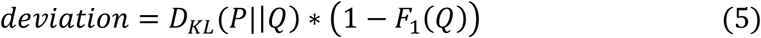

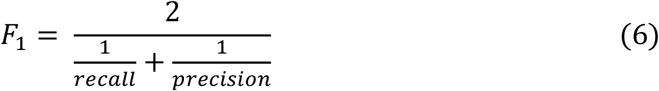

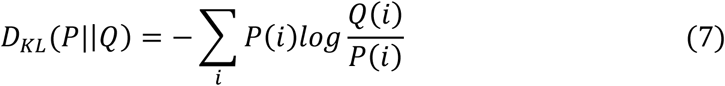 F1 score is the harmonic mean of precision and recall, which optimally blend both of precision and recall metrics and can comprehensively characterize the performance of methods. KLD could quantify the diverges between two probability distributions (Equation 7), the D_KL_(*P*‖*Q*) represents the divergence from *Q* to *P*. In this study, *Q* is the distribution of CDR3 repertoires retrieved from each tool, and the P represents the distribution of “true” CDR3s in simulation data. Combined the two measures could compare the precision of tools when their detected CDR3 repertoires have similar distributions.
3. *Computational resource*: Measured by time cost and memory usage. Time cost is the sum of user time and system time from built-in *time* command, which can reflect the actual consumption of computing power in terms of the number of thread/process.

---

**Figure S1.**
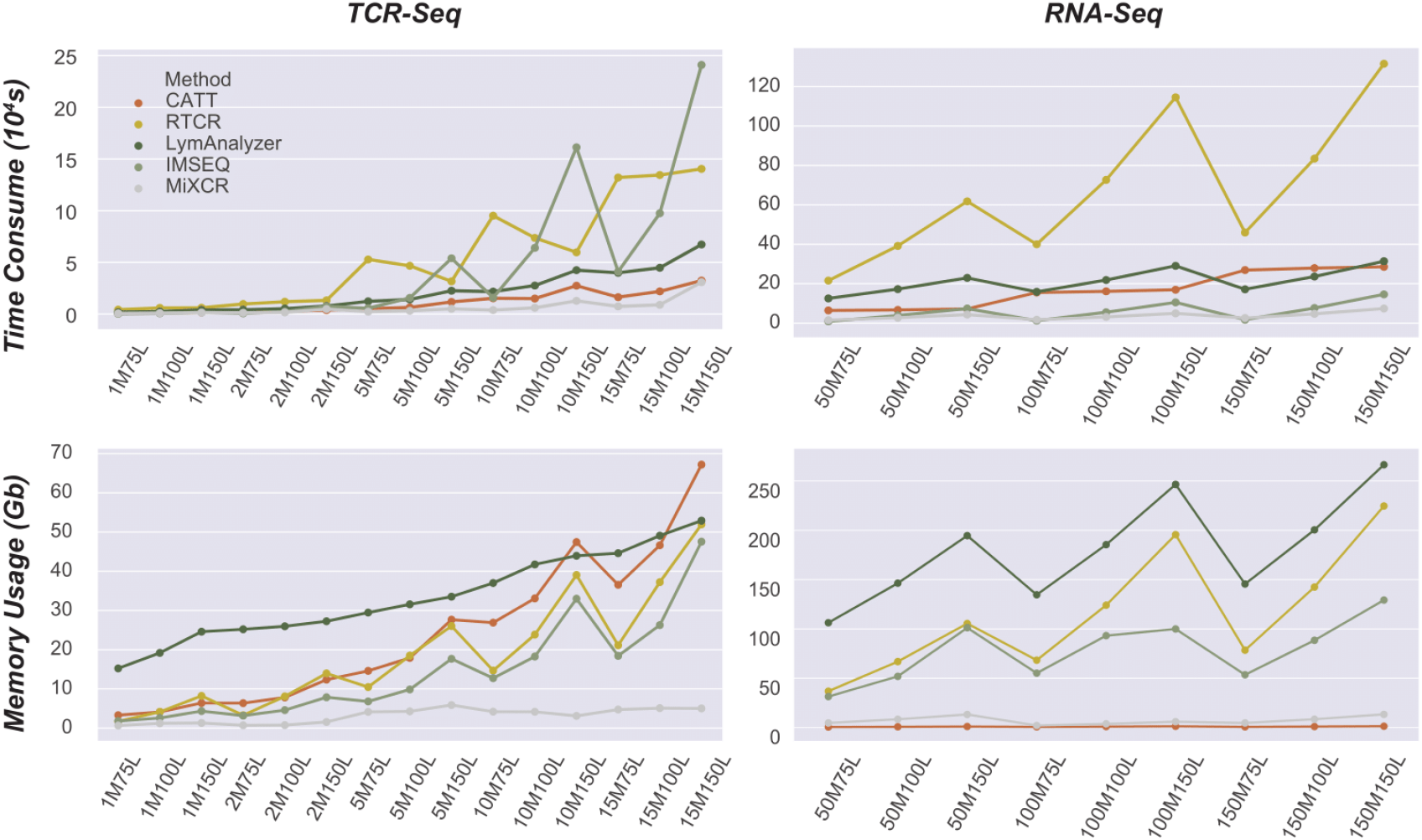
The memory usage and time consume of each tool on in-silico TCR-Seq datasets and RNA-Seq datasets. The time consume was measured by the sum of system time and user time.

**Figure S2.**
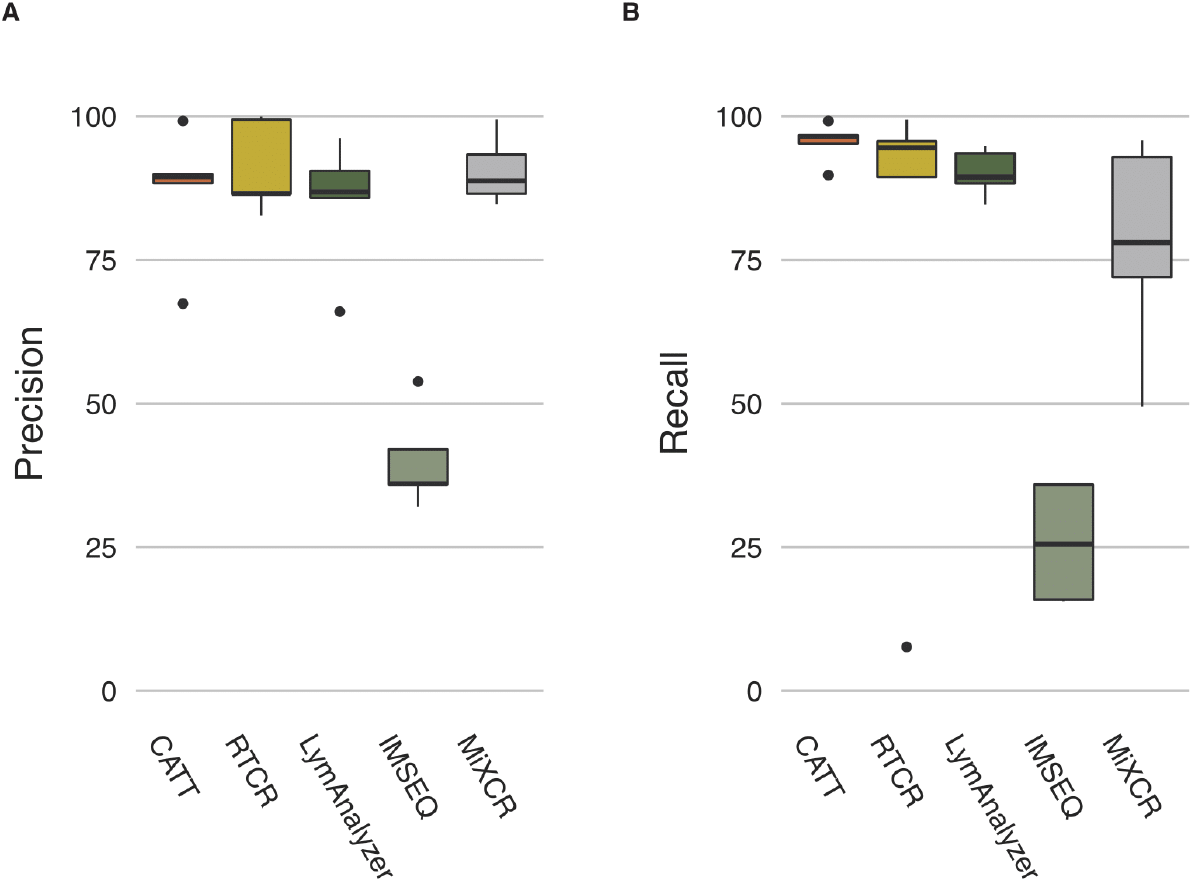
The precision (A) and recall(B) for each tool on scTCR-Seq data

**Figure S3.**
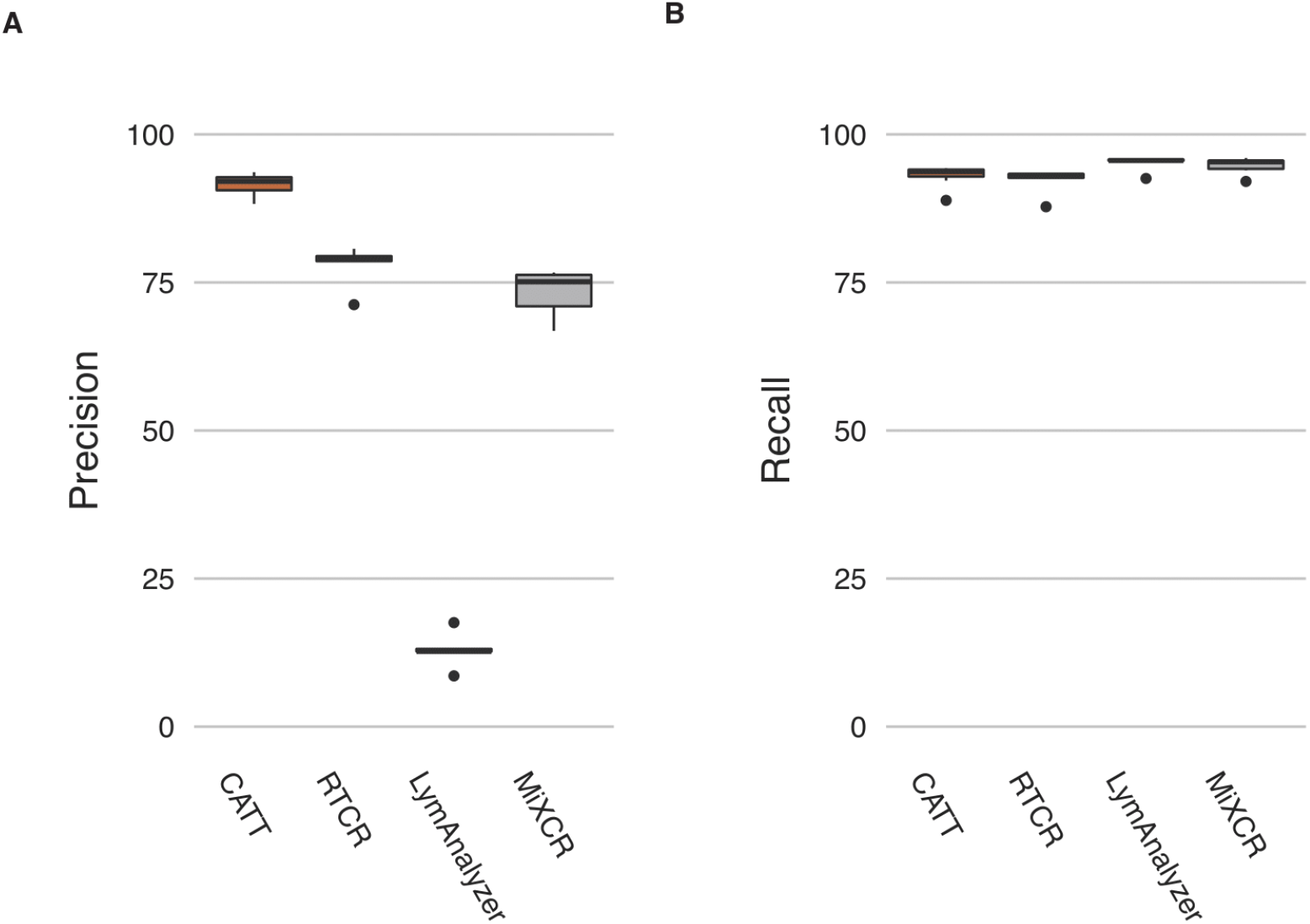
The precision (A) and recall (B) of each tool on TCR-Seq dataset.

**Figure S4.**
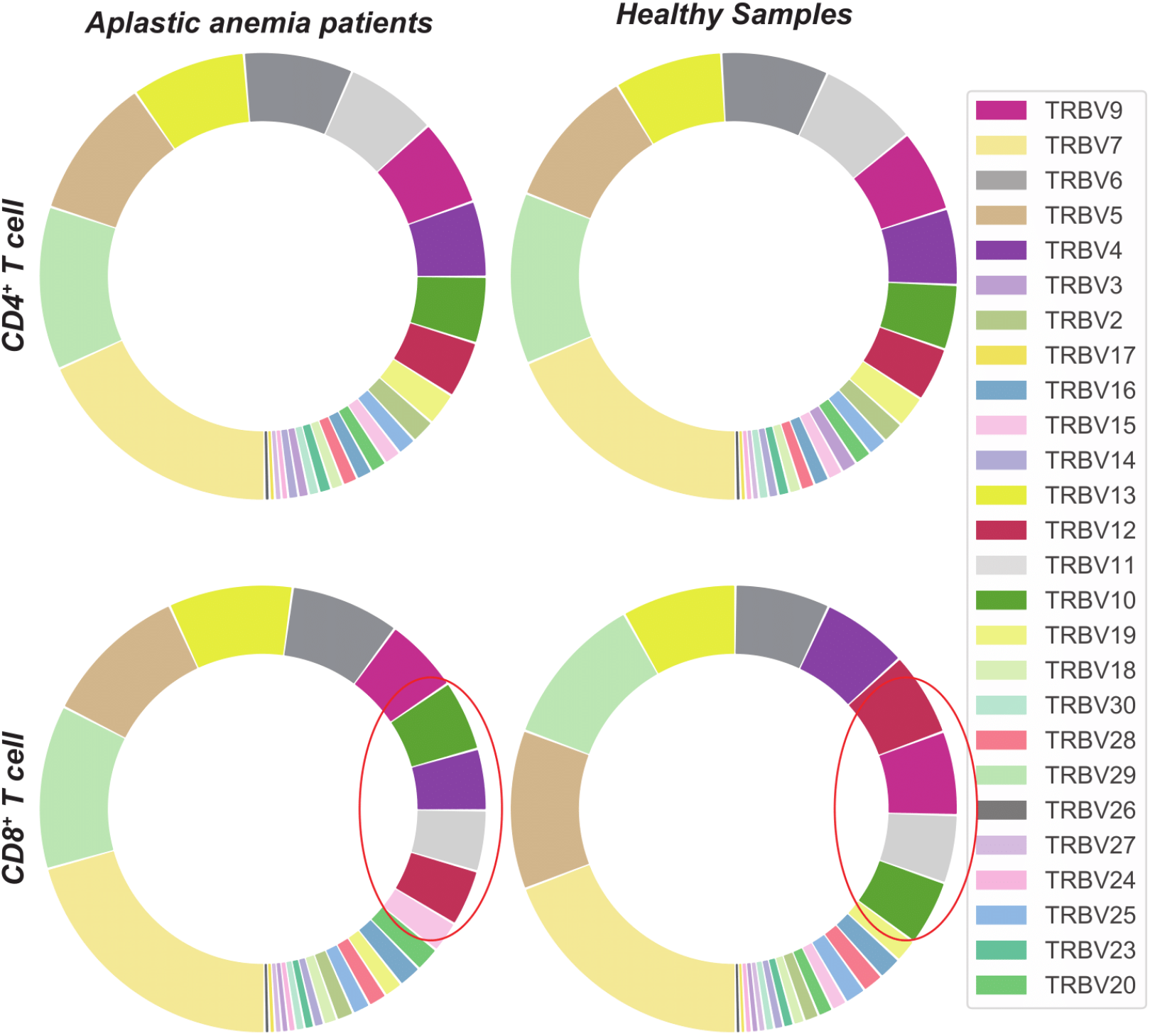
The TRBV gene usage in CD8+ T cells and CD4+ T cells of healthy donors and aplastic anemia patients.

## Acknowledgements

We thank Dr. Zhichao Chen from Wuhan Union Hospital for helpful discussions. We thank the Zi-Yi Song for providing the inspiration of the tool name.

## Code availability

The source codes for CATT are freely available at http://bioinfo.life.hust.edu.cn/CATT and GitHub (https://github.com/GuoBioinfoLab/CATT). CATT is written in Python 3/Julia and platform independent. It is released under the GNU AFFERO General Public License v3.0.

